# Planar polarized force propagation integrates cell behavior with tissue shaping during convergent extension

**DOI:** 10.1101/2022.11.08.515701

**Authors:** Shinuo Weng, John B. Wallingford

## Abstract

Convergent extension (CE) is an evolutionarily conserved developmental process that elongates tissues and organs via collective cell movements known as cell intercalation. Here, we sought to understand the mechanisms connecting cell behaviors and tissue shaping. We focus on an often-overlooked aspect of cell intercalation, the resolution of 4-cell rosettes. Our data reveal that polarized cellular forces are involved in a timely rosette resolution, which in turn, enables propagation of such cellular forces, facilitating the propagation of tissue-scale CE. Conversely, delayed rosette resolution leads to a subtle but significant change of tissue-wide cell packing and exerts a profound impact by blocking force propagation, resulting in CE propagation defects. Our findings propose a collaborative nature of local cell intercalations in propagating tissue-wide CE. It unveils a multiscale biomechanical synergy underpinning the cellular mechanisms that orchestrate tissue morphogenesis during CE.

**Highlights:**

- 4-cell rosette is resolved by a two-step process: t-junction extension, then rotation.
- Delayed t-junction rotation significantly impacts the tissue-wide cell packing configuration.
- Timely resolved 4-cell rosettes enable polarized force coupling and propagation both *in silico* and *in vivo*.
- Polarized force propagation is involved in the propagation of tissue shaping.

## Introduction

Convergent extension (CE) is an evolutionarily conserved morphogenetic process crucial for the elongation of the body axis in almost all animals and for the development of multiple organ systems during embryogenesis (Tada and Heisenberg, 2012, Shindo, 2018, Sutherland et al., 2020). CE involves intricate cellular behaviors, including planar cell polarity establishment, polarized actomyosin activity, and directed cell intercalations (as shown in **Fig. 1A**). These cell behaviors act collectively to narrow a tissue in one direction while elongating it in another direction (Devenport, 2016, Huebner and Wallingford, 2018, Shindo, 2018).

**Figure 1.**
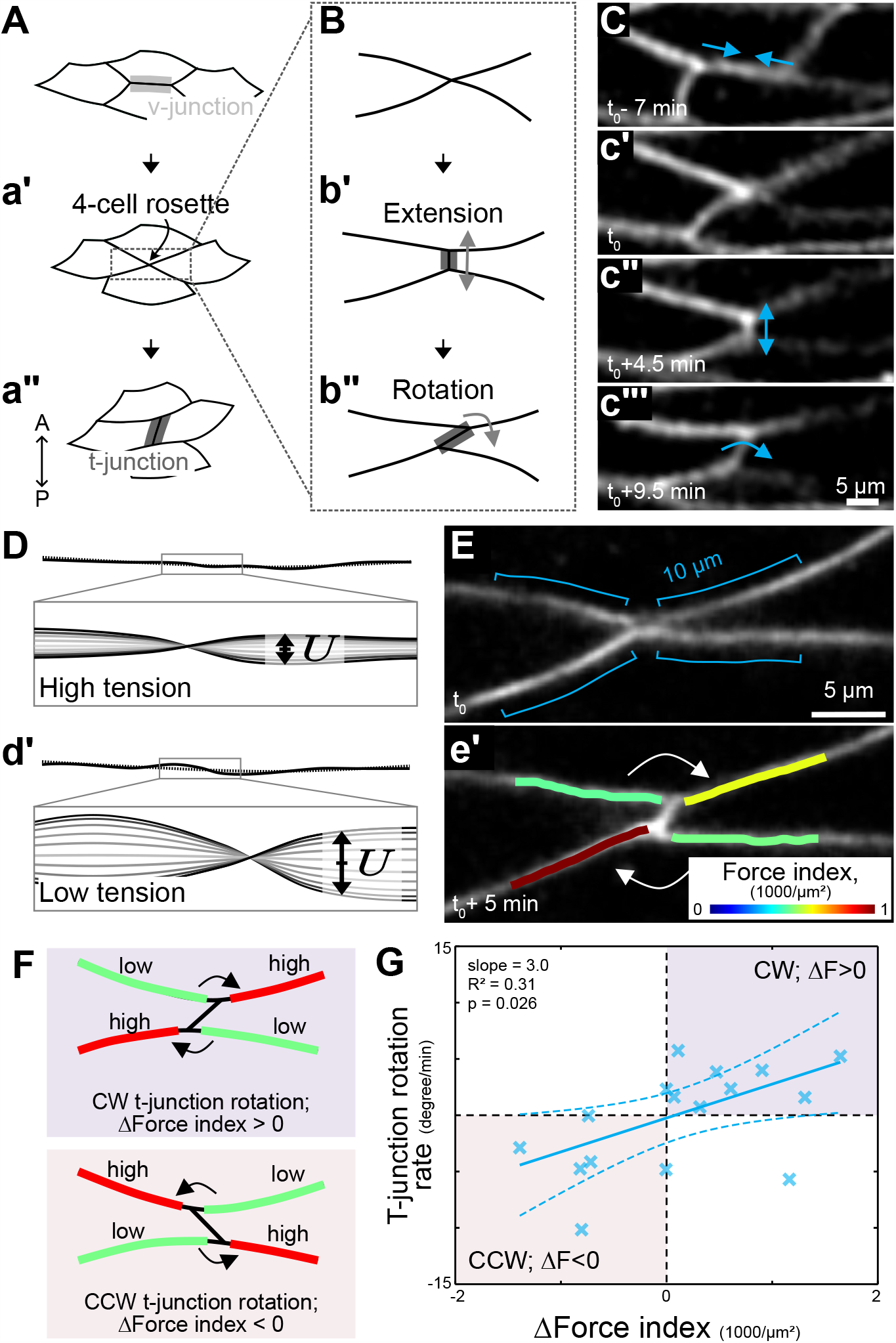
4-cell rosette resolves by t-junction extension and then rotation. **(A)** Schematic showing a 2-step process of cell intercalation: mediolateral cells shorten v-junctions (light gray) while forming a 4-cell rosette (**a’**), followed by nascent t-junction (dark gray) formation as cells continue to move (**a”**). **(B)** Schematic showing a 2-step resolution of 4-cell rosettes: nascent t-junctions extend anteroposteriorly first(**b’**) and rotate afterward (**b”).** **(C)** Still images from representative movies showing the entire process of cell intercalation. **(D)** Schematic correlating transverse fluctuation, *U*, of a junction with its junctional tension. High junctional tension correlates with low transverse fluctuation, and vice versa. See **Method.** **(E)** Representative images of t-junction dynamics and tension analysis on connected v-junctions. Transverse fluctuation, *U*, on the adjacent 10-μm region of each connected v-junction during the first 5-min of the t-junction formation was measured. The force index, 1/⟨|*U*|^2^⟩, was quantified and displayed in a color-coded manner as an overlay on top of the junction image. **(F)** Schematic depicting that diagonal v-junctions exert planar polarized tension to control the rotation of connected t-junctions. Top panel, if the tension on the upper right and lower left v-junctions are higher than the tension on the other diagonal pair, the connected t-junction will rotate in the clockwise direction. This scenario corresponds to the data points in the first quadrant in (**G**). Bottom panel, if the tension on the upper left and lower right v-junctions are higher, the t-junction will rotate counterclockwise. This corresponds to data points in the third quadrant in (**G**). **(G)** Plot showing t-junction rotation rate versus the force index difference on diagonal v-junctions (Δ*Force Index*). A positive value of Δ*Force Index* indicates higher tension on the upper right and lower left v-junctions, and vice versa. Cross markers represent data points, the solid line is the linear fitting, and the dashed lines are upper and lower boundaries with 95% confidence. N = 16.

A major outstanding question relates to the mechanisms by which individual cell behaviors are integrated to achieve shape change at the level of whole tissues. An attractive candidate is the patterned spatial arrangement of cells, known as cell packing. At larger scales, packing serves as a remarkable predictor of collective cell motility (Staple et al., 2010, Bi et al., 2015, Wang et al., 2020) and tissue material properties (Petridou et al., 2021). At the same time, packing is known to be influenced by cellular processes involving neighbor exchanges (Gibson et al., 2006, Wang et al., 2020),

Neighbor exchange by cell intercalation is the key driver of CE and it comprises two pivotal steps: the formation of a multi-cell rosette, where so-called v-junctions connecting anteroposterior (AP) cell neighbors shorten to bring the mediolateral (ML) cell neighbors into contact (**Fig. 1A-a’**), and the resolution of the rosette, in which a nascent t-junction forms and grows between the neighbors (**Fig. 1a’ -a”**). While most studies of CE have focused on the first step of cell intercalation, it is the far less-explored second step, resolution, that actually elongates the tissue, as evidenced by studies on *Drosophila* germ band extension (Collinet et al., 2015, Yu and Fernandez-Gonzalez, 2016). Nevertheless, our understanding of rosette resolution and its role in tissue shaping, especially in vertebrate models such as *Xenopus*, remains significantly limited.

Here, we combined embryology, cell biology, and novel tools for the non-invasive assessment of mechanical forces across different developmental scales to link the resolution of cell intercalation, cell packing configurations, and tissue scale CE in *Xenopus* notochord. Our findings illuminate the role of timely t-junction rotation, an overlooked process at the end of cell intercalation, in maintaining normal cell packing that facilitates polarized propagation of cellular forces and thus, effective tissue-level CE. Our data provide multiscale mechanical insights into the propagation of tissue shaping through intricate and collaborative cell behaviors.

## Results

### Nascent t-junctions resolve 4-cell rosettes by extension and then rotation

We first characterized the resolution of 4-cell rosettes during CE in the *Xenopus* notochord by live imaging of a membrane marker (**Fig.1 B,C**). Curiously, and contrary to the single-step process depicted in **Fig. 1a”**, our data unveiled a two-step process in rosette resolution. Initially, nascent t-junctions extended perpendicularly to the connected v-junctions, as shown in **Fig. 1b’, c”**. This extension occurred at an approximate rate of 1 μm/min, spanning a duration of roughly 3 minutes with the nascent t-junction reaching a length of about 3 μm. Subsequently, in a second phase, the t-junction displayed an acute rotational movement while continuing to elongate (**Fig. 1b”, c’”**), proceeding at a rate of ∼6 º/min. This second step in nascent t-junction growth seems not to have been described previously and prompted us to ask what cellular mechanisms drive this process.

### T-junction rotation involves differential high tension on the connected junctions

Previous work by our lab and others has shown that mediolaterally aligned v-junctions bear greater tension than the relatively anteroposterior aligned t-junctions (Shindo and Wallingford, 2014, Weng et al., 2023). This increased tension on v-junctions drives v-junction shortening in the first half of cell intercalation, so we reasoned that the rotation of nascent t-junctions may then be driven by tension from their adjoining ML-aligned junctions.

To test this idea, we applied a newly developed image-based, non-invasive method named Tension by Transverse Fluctuation, or “TFlux”, to assess junctional tension *in vivo* (see **Method** for details) (Weng et al., 2023). Briefly, TFlux measures the transverse fluctuation of cell membrane along cell-cell interfaces over time, and the force index is inversely proportional to the mean square transverse fluctuation (**Fig. 1D**). This image-based analysis enabled us to directly link t-junction growth dynamics with tensions from its adjoining ML aligned junctions (**Fig. 1E**).

We imagined the process of t-junction rotation as a result of a 4-way tug of war. In this model, a new t-junction rotates clockwise when tensions on the upper right and lower left adjoining junctions are higher than tensions on the upper left and lower right ones (**Fig. 1F**, top; example in **Fig. 1e’**). Conversely, a new t-junction rotates counterclockwise when this differential mechanical pattern is flipped (**Fig. 1F**, lower).

To quantify this model, we introduce the term Δ*Force Index*, representing the tension difference between the diagonal pairs of connected junctions, such that a positive value indicates higher tension on the upper right and lower left v-junctions, and a negative value, the reverse. We plotted Δ*Force Index* against the overall t-junction rotation rate and we found that 15 out of 16 data points were in the first and third quadrants (**Fig. 1G**), consistent with our hypothesis illustrated in **Fig. 1F**. These data suggested that planar polarization of tensions on ML-aligned junctions determines the rotation direction of the conjoined t-junction.

### Lagging rosette resolution alters tissue-wide cell packing configuration

We recently found that knockdown of the catenin Arvcf subtly decreases cell cortex tension during CE (Huebner et al., 2022), suggesting that it might be involved in t-junction rotation. Indeed, live imaging of Arvcf-deficient cells during rosette resolution revealed that nascent t-junctions formed (**Fig. 2A**), but both their extension and rotation displayed significantly altered dynamics (**Fig. 2B-E**). Loss of Arvcf led to a 20% reduction in the extension rate compared to wildtype (WT) cells (**Fig. 2B**) and prolonged the extension period by over 250% (**Fig. 2C**). As a result, t-junctions were notably longer at the onset of rotation (**Fig. 2D**). Furthermore, the rotation rate exhibited a 60% reduction (**Fig. 2E**), suggesting that it is the impeded t-junction rotation that delays the transition from extension to rotation.

**Figure 2.**
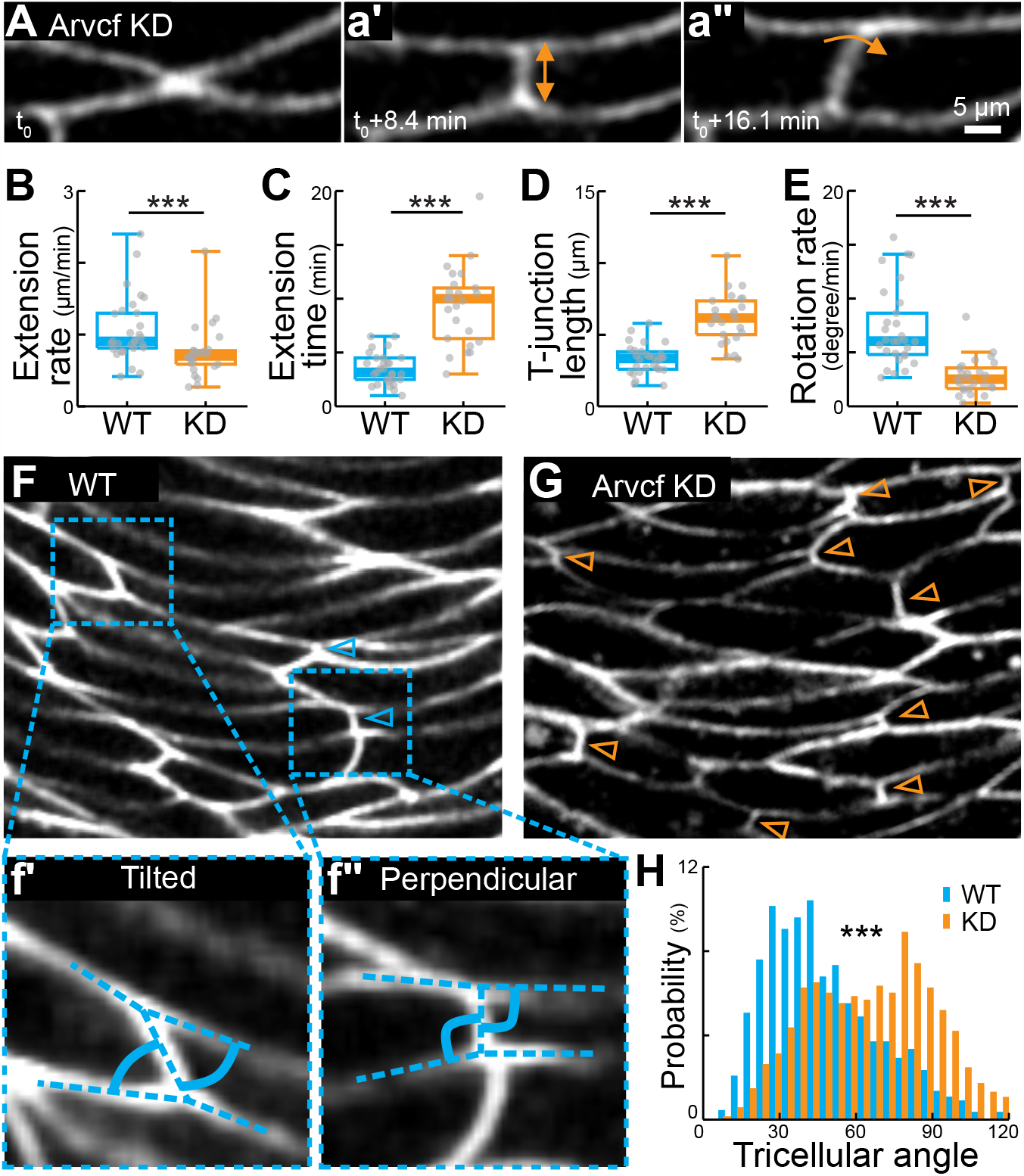
Cell packing configuration is finely tuned by the t-junction rotation rate. (**A**) Still images from representative movies illustrating t-junction formation in Arvcf deficient cells. Left panel, a t-junction starts to form when the ML cells meet. Middle panels, t-junction extends along the AP axis. Right panels, t-junction rotates at the end of the extension phase. (**B-E**) Quantification of a nascent t-junction dynamics, including the extension rate (**B**), extension time (**C**), junction length before rotation (**D**), and the rotation rate (**E**) in WT and Arvcf KD explants. N = 32 for t-junctions from WT explants and N = 26 from Arvcf KD explants. Explants were collected from at least three replicates. p values were calculated using Wilcoxon rank sum test (A.K.A. Mann-Whitney U test). ***p < 0.0005. (**F-G**) Representative images of cell packing configurations in WT (**F**) and Arvcf KD (**G**) explants with membrane labeling. Insets (**f’, f”**) are representative images showing the angle between t- and connected v-junctions (“tricellular angles”; solid lines). Arrowheads mark the so-called AP aligned t-junctions perpendicular to the surrounding v-junctions. (**H**) Distribution of the tricellular angle, defined as the smaller angle between t- and connected v-junctions for each cell (examples in **f’** and **f”**). Data points were pooled from 11 explants for each condition from at least three replicates and approximately 50 cells from each explant. p value was calculated using Kolmogorov–Smirnov test. ***p < 0.0005.

Interestingly, longer extension and slower rotation translated into noticeable differences in cell packing configurations between WT and Arvcf KD explants. At the population level, t-junctions in control explants typically displayed a tilted orientation relative to their adjoining junctions (**Fig. 2F,2f’**), with only a limited subset of them being perpendicular (i.e., strongly aligned in the AP axis at the cellular scale; **Fig. 2f”**, arrowheads). To quantify this phenotype, we measured the smaller angle between each t-junction and its connected junctions (referred to as “tricellular angles”, **Fig. 2f’, f”**, solid lines). The distribution of tricellular angles peaked at around 40º in control explants (**Fig. 2H**, blue), consistent with the tilted pattern described above.

Conversely, Arvcf depletion led to an increased prevalence of AP-aligned t-junctions throughout the population (**Fig. 2G**, arrowheads). The tricellular angle measure further confirmed this observation: with a dominant peak around 90º (**Fig. 2H, orange**). Coupled with the observations in **Fig. 2A-E**, our data suggested that Arvcf deficiency prolonged the resolution of 4-cell rosettes by delaying the extension-to-rotation transition in new t-junctions, and this, in turn, led ultimately to an overall shift in cell packing configuration characterized by an excess of AP-aligned junctions within the tissue.

### Cell packing configuration facilitates planar polarized force propagation *in silico*

Embryos lacking Arvcf display severe defects in axis elongation despite relatively subtle defects in intercalation (Fang et al., 2004, Weng et al., 2022, Huebner et al., 2022), leading us to ask if the distinct cell packing configurations we observed might affect CE. We considered mechanical principles, akin for example to how the pattern of beams and posts forming a truss bridge influence the distribution of loading. We first explored this idea by creating computational toy models to analyze relative force distributions in a tissue using a force balance-based inference technique (**Fig. 3A**)(Brodland et al., 2014). In an initial experiment, we tested models that represented extreme cell packing configurations, deliberately exaggerating features observed in Control and Arvcf KD embryos (**Fig. 3B-C**). The first model, termed *Hexagon*, employed cells with hexagonal shapes and tilted t-junctions (**Fig. 3B**). The second model, termed *Brick-wall*, featured rectangular cells with all t-junctions perpendicular to the neighboring junctions (**Fig. 3C**).

**Figure 3.**
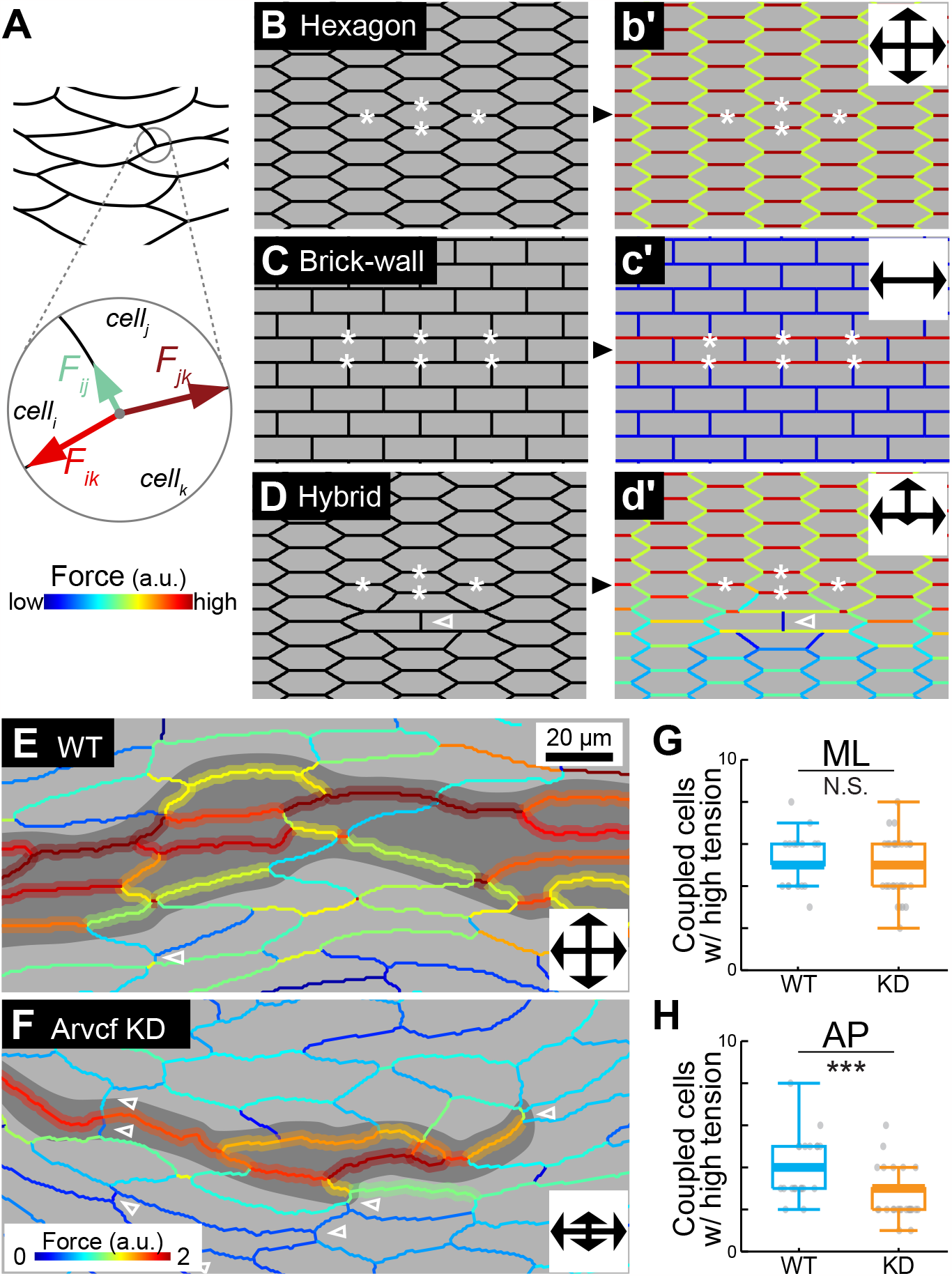
Resolution of 4-cell rosettes is required for polarized force propagation. (**A**) Schematic explaining the cellular force interference technique, illustrating force balance at each tricellular vertex. (**B-D**) Force propagation simulation in toy models with different cell packing configurations. Left panel, 3 representative cell packing configuration - A *Hexagon* model (**B**), *Brick-wall* model (**C**), and *Hybrid* model (**D**). Right panel, force distribution across the tissue model upon force stimulation in the center, indicating relative tension on each junction with color code. Dark red is the highest and blue is the lowest. Insets show the overall force propagation patten. (**E, F**) Force distribution estimated using the cellular force inference in WT (**E**) and Arvcf KD (**F**) explants. Dark gray shading highlights regions with v-junctions having the maximum force (dark red), along with their adjacent v-junctions holding forces greater than the average (warm color). Arrowheads mark the t-junctions perpendicular to the apposed v-junctions. (**G, H**) Quantification of force coupling in the ML (**G**) and AP (**H**) directions in WT and Arvcf KD explants. The number of adjacent cells having forces on v-junctions above average were quantified as the number of cells with force coupled. Data pooled from 20 WT explants and 17 Arvcf KD explants from at least three replicates. p values were calculated using Wilcoxon rank sum test (A.K.A. Mann-Whitney U test). ***p < 0.0005; NS, not significant.

We then computationally applied high tension to specific v-junctions in the center of the tissue (marked by asterisks in **Fig. 3B, C**) to mimic the ML converging forces of cell intercalation. The results revealed striking differences in force propagation between the two models, as indicated by force heatmaps in **Fig. 3b’** and **c’** (index shown in **Fig. 3A**). In the *Hexagon* tissue, forces applied in the center propagated in both ML and AP directions, creating high tensions across v- and t-junctions throughout the tissue (**Fig. 3b’**, red/yellow). In contrast, the *Brick-wall* tissue allowed only ML force propagation, creating two ML-aligned chains of high tension (**Fig. 3c’**, red); neither t-nor v-junctions above or below the applied forces displayed any elevated tension (**Fig. 3c’**, blue).

As an additional test, we introduced a *Hybrid* tissue model, incorporating a single AP-aligned t-junction in the *Hexagon* tissue (**Fig. 3D**, arrowhead). Forces applied in this hybrid tissue (asterisks) propagated in the ML direction and also towards the anterior (up) (**Fig. 3d’**). However, force propagation towards the posterior (down) exhibited an immediate drop at the inserted AP-aligned t-junction (**Fig. 3d’**). These preliminary model tests suggested that AP-aligned t-junctions can efficiently block AP force propagation at the tissue level.

### Aberrant cell packing configuration disrupts planar polarized force coupling during CE *in vivo*

We proceeded to validate our force propagation model *in vivo* by evaluating force distribution in tissue explants during CE using the same cellular force inference tool (Brodland et al., 2014). We found that force distribution was highly heterogeneous across cells within WT explants, with regions of significantly higher and lower tension (**Fig. 3E**). Interestingly, however, the regions of high tension consistently spanned four to five cell diameters in both the AP and ML directions (**Fig. 3E**, highlighted; **Fig. 3G, H**, blue), suggesting robust force coupling and propagation in both directions.

By contrast, the heterogenous force distribution in Arvcf KD explants displayed a different pattern. High-tension regions manifested as chains, coupled in the ML direction but not in the AP direction (**Fig. 3F; Fig. 3G, H**, orange). This pattern resembled the force propagation pattern witnessed in the *Brick-wall* toy model (**Fig. 3c’**). Notably, a considerable number of AP-aligned t-junctions surrounded the high-tension region in Arvcf KD explants (**Fig. 3F**, arrowheads). These observations suggest that the excessive AP-aligned t-junctions in the Arvcf deficient tissue effectively block the AP propagation of forces from local cell intercalations.

### Defects in rosette resolution and cell packing disrupt the orderly propagation of convergent extension in the notochord

Lastly, we ask if delayed rosette resolution, altered cell packing, and hindered force propagation at relatively smaller scales translate into differences in tissue shaping at larger, tissue and embryo scales. We conducted live-imaging of notochord explants undergoing CE over a 2-hour period. At the beginning, WT explants displayed a trapezoidal shape (**Fig. 4A**), consistent with anterior-to-posterior propagation of CE cell behaviors including cell elongation and intercalation, a phenomenon reported decades ago (Shih and Keller, 1992). Over the two-hour imaging, we observed substantial AP extension and ML convergence across the entire AP axis (**Fig. 4A**). Convergence was quantified at three positions along the AP axis (**Fig. 4C**, left), showing consistent convergence thoroughly during this time (**Fig. 4C**, blue).

**Figure 4.**
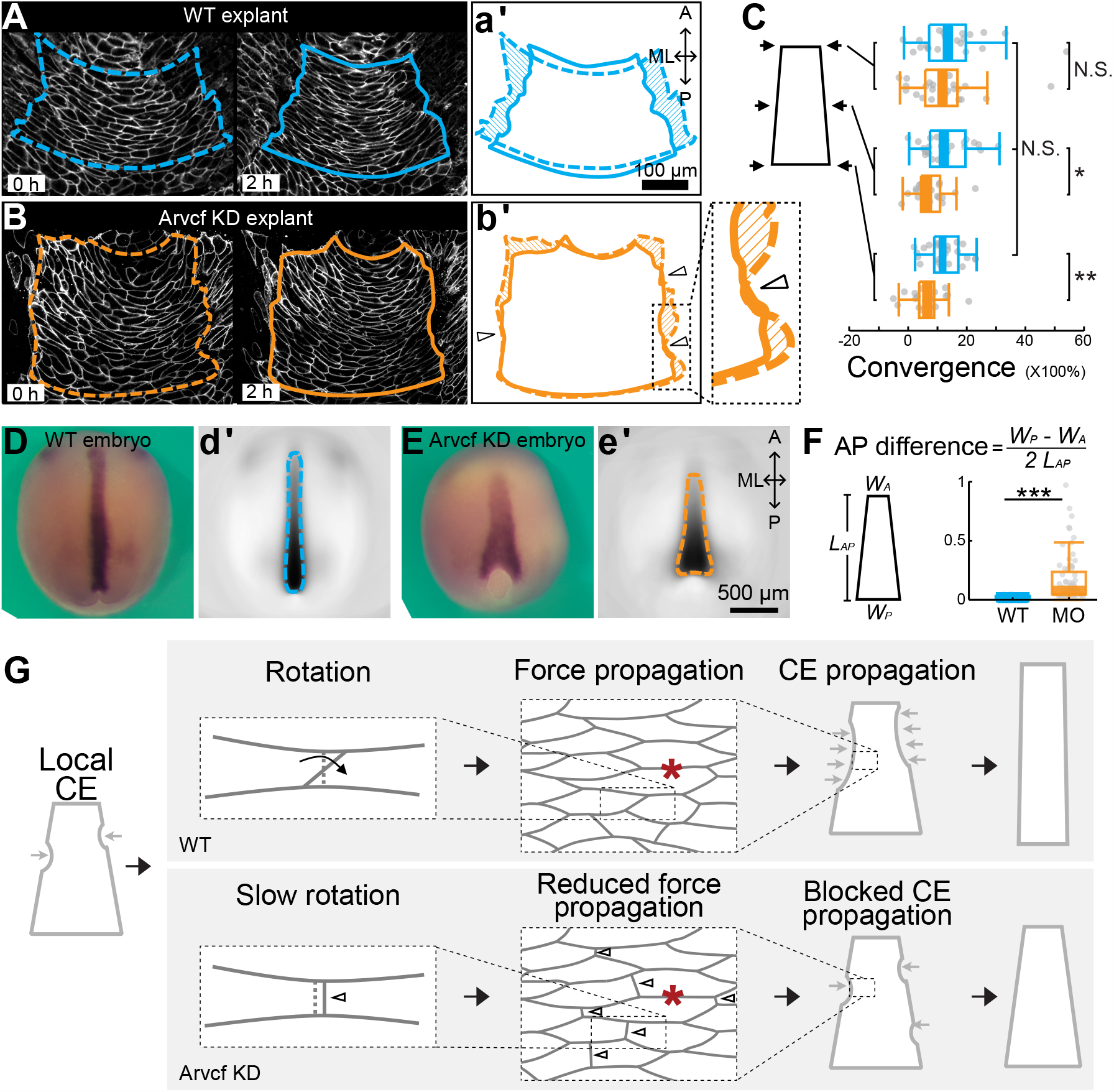
Propagation of tissue-scale CE is required for efficient tissue shaping. (**A, B**) Convergent extension of Wildtype (**A**) and Arvcf KD (**B**) explants during a two-hour incubation. Dashed lines mark tissue region an equivalent stage of 13. Solid lines indicate the same region post-incubation. Two outlines were stacked together to show the shape change, highlighted with hatching shades (**a’, b’**). Arrowheads mark the regions with interrupted convergences along the AP axis. (**C**) Quantification of ML convergence at the anterior, middle, and posterior ends as illustrated in the left panel. N = 24 for WT explants (blue) and N = 23 for Arvcf deficient explants (orange) from at least three replicates. p values were calculated using Wilcoxon rank sum test (A.K.A. Mann-Whitney U test). *p < 0.05; **p < 0.005; NS, not significant. (**D, E**) Wildtype (**D**) and Arvcf KD (**E**) embryos at stage 14 stained by *in situ* hybridization for the notochord-specific probe *Xnot*. Left, representative images of embryos. Right, stacked grayscale images of over 70 embryos for each condition. Dashed lines outline the average shape of a notochord with 50% coverage. **(F)** Quantification of normalized width difference between posterior and anterior notochords. p values were calculated using Wilcoxon rank sum test (A.K.A. Mann-Whitney U test). ***p < 0.0005. **(G)** Multiscale biomechanical model underpinning tissue-scale CE propagation. Cell intercalation causes local CE. In WT, timely t-junction rotation resolves 4-cell rosettes, enabling forces from other cell intercalation events (asterisks) to propagate in the AP direction, promoting CE propagation. Arvcf loss impedes t-junction rotation, leading to excessive AP aligned t-junctions that block AP force propagation, hindering CE propagation.

Interestingly, when we quantified the ML convergence in Arvcf deficient explants, we observed relatively normal convergence at the anterior end of the tissue, but significantly reduced convergence toward the posterior (**Fig. 4C**), leading to a reduced overall ML convergence (**Fig. 4B**). However, the anterior-to-posterior propagation of CE cell behaviors remained largely unaffected, as we consistently observed elongated and polarized cells with active cell intercalation at all positions (data not shown), leading to multiple instances of localized convergence in the middle-to-posterior region of Arvcf-deficient explant (**Fig. 4b’**). Intriguingly, these localized CE regions were surrounded by areas where convergence was minimal to nonexistent (**Fig. 4b’**, indicated by arrowheads), suggesting that localized tissue-scale convergence fails to propagate effectively along the AP axis.

Finally, we asked how this tissue-scale CE propagation defect impacted notochord shape in the embryo. Using *in situ* hybridization for the notochord-specific marker *Xnot* (von Dassow et al., 1993), we found that the notochord in control embryos at early neurulation (NF stage 14) had reorganized into a slim rectangle (**Fig. 4D**), with little difference in tissue width between its anterior and posterior (**Fig. 4F**, blue). By contrast, the shorter and wider notochords of Arvcf KD embryos (Huebner et al., 2022) displayed notochords that had narrowed anteriorly but remained significantly wider at the posterior (**Fig.4E; 4F**, orange), consistent with the observations in tissue explants (**Fig. 4C**). These data collectively argue that timely resolution of 4-cell rosettes after cell intercalation are essential for normal cell packing which in turn is essential for normal force propagation across the tissue and ultimately for the normal patterning of tissue-scale convergent extension (**Fig. 4G**).

## Discussion

Our overarching goal was to unravel the mechanisms connecting microscale and macroscale behaviors to achieve efficient tissue-scale morphogenesis. By employing a combination of live-imaging, image-based mechanical measurements, and computational modeling, we have pinpointed the nascent t-junction rotation at the end of each cell intercalation event as the key to bridge CE behaviors across different scale.

To understand its implications, we delved into the realm of cell packing configuration. This concept has been linked to collective cell motility as suggested by various vertex models designed for epithelia (Bi et al., 2015, Wang et al., 2020). However, the currently considered parameters such as cell shape index and cell alignment index, do not exhibit significant changes associated with t-junction rotation in *Xenopus* notochord. This highlights the pressing need for new models that incorporate additional factors.

Our analysis of cell packing underscores the significance of force propagation capability in facilitating productive tissue shaping. Specifically, we demonstrated that the propagation of cellular forces along the AP axis depends on the cell packing configuration, which is finely tuned by the orientation of t-junctions (**Fig. 2**,**3**). This force propagation is further implicated in the propagation of tissue-scale CE (**Fig. 4**). Interestingly, a similar concept of a mechano-dependent morphogenetic wave has been described in *Drosophila* endoderm invagination (Bailles et al., 2019). It worth exploring whether a tissue-scale wave of Rho1/MyoII activation, in addition to the known calcium wave (Wallingford et al., 2001), exists in the *Xenopus* notochord, and how these waves correlate with cell packing configuration.

Finally, we explored the cellular mechanisms driving t-junction rotation and demonstrated that high tension on the ML aligned junctions, aside from its well documented roles on v-junction shortening, is involved in t-junction rotation (**Fig. 1E-G**). The pulsatile nature of these asynchronized high tension events (Shindo et al., 2019) generates a polarized force pattern necessary for rapid t-junction rotation. Notably, a recent vertex model of CE has presented a scenario in which cellular forces are synchronized across the tissue, leading to creation of perfectly AP aligned t-junctions (although not discussed in the original work). Interestingly, this hypothetical scenario brings tissue-level CE to a complete halt after just one round of cell intercalation (Shindo et al., 2019), an exaggerated case akin to **Fig. 4B**. Collectively, our data suggest a multifaceted biomechanical linkage between individual cell intercalation events and efficient tissue shaping.

## Acknowledgements

We thank Dan Dickinson for his helpful comments and critical reading. This work was supported by R01HD099191.

## Author contributions

S.W. and J.B.W. conceptualized the project and wrote the manuscript. S.W. designed and conducted the experiments, performed analyses, and developed the model.

## Methods

### *Xenopus* embryo manipulations

Ovulation was induced by injecting adult female Xenopus laevis with 600 units of human chorionic gonadotropin (HCG, MERCK Animal Health) and animals were kept at 16 dc overnight. Eggs were acquired the following day by squeezing the ovulating females and eggs were fertilized *in vitro*. Eggs were dejellied in 2.5% cysteine (pH 7.9) 1.5 hours after fertilization and reared in 1/3x Marc’s modified Ringer’s (MMR) solution. For micro-injection, embryos were placed in 2% ficoll in 1/3x MMR during injection and washed in 1/3x MMR 30 min after injection. Embryos were injected in the dorsal blastomeres at the 4-cell stage targeting the C1 cell at 32-cell stage and presumptive notochord. Keller explants were dissected at stage 10.25 in Steinberg’s solution using hair tools.

### Plasmids and Morpholinos

The Arvcf morpholino has been previously described (5’-ACACTGGCAGACCTGAGCCTATGGC-3’ (Fang et al., 2004) and was ordered from Gene Tools, LLC. Membrane-BFP plasmid was made in pCS2.

### mRNA and morpholino microinjections

Capped mRNAs were generated using the ThermoFisher SP6 mMessage mMachine kit (Catalog number: AM1340). Membrane-BFP mRNAs were injected at 75 pg per blastomere.

### *In situ* hybridization

Whole-mount *in situ* hybridization was performed as descried previously using a DIG-labeled single-strand RNA probe against a partial sequence of *Xnot* (Monsoro-Burq, 2007, Sive et al., 2000). This antisense probe has been well characterized, which shows *Xnot* expression in the prenotochordal region about the dorsal lip at stage 10.5 and along the dorsal midline exclusive to the notochord up to stage 16 (von Dassow et al., 1993). Bright field images were captured with a Zeiss Axio Zoom. V16 stereo microscope with Carl Zeiss Axiocam HRc color microscope camera or a Leica stereo microscope MDG41 with AmScope microscope digital camera WF200.

### Imaging *Xenopus* explants

Explants were submerged in Steinberg’s solutions and cultured on glass coverslips coated with Fibronectin (Sigma-Aldrich, F1141) at 5 μg/cm^2^. All images of membrane labeling (Membrane-BFP) were taken on a Nikon A1R, and at the focal plane 5 μm deep into the explant. For tracking tissue-scale CE, the first image was taken after a 3-hour incubation at room temperature. A second image was taken another two hours later. All other assays were started after a 5-hour incubation of the explant. For tracking the dynamic of t-junction formation, we set the standard confocal time-lapse imaging at an interval of 5 sec. For the force inference by transverse fluctuation, we set the time interval of 1 sec.

### Tension by Transverse Fluctuation (TFlux)

TFlux measurement is based on the idea that the transverse fluctuation of a junction is inversely proportional to tension on the junction (**Fig. 1D**) (Weng et al., 2023). To track the junction movement, we labeled the membrane by injecting embryos at the 4-cell stage in both dorsal blastomeres with Mem-BFP mRNA. Keller explants were then dissected from early gastrula embryos, mounted on fibronectin coated coverslips, incubated in the Steinberg’s solution at room temperature for five hours, and then live-imaged on a Nikon A1R confocal microscope for 5 min. To capture small transverse fluctuations along a junction, we used time interval of 1 sec and pixel size of 0.20 μm. To quantify the transverse fluctuations, time-lapse movies were imported into Ilastik® and apposing cells were first segmented using Ilastik® pixel classification and Ilastik® Carving. 3D (xyt) meshes of the connected cells were then imported to a customized MATLAB script to detect the transverse fluctuation over time. Briefly, we first defined a base-line position of the junction at each time point to account for the overall movement of the cells and the junction. To do so, we performed a moving average over 2 μm along the junction length at each time point, and then another moving average over 20 sec. Transverse fluctuation was then measured at each time point as the distance from the original junction position to the base line. The overall junction tension, referred to as the force index, is defined as the reciprocal of the mean square transverse fluctuation of the entire junction of interest over the 5-min of observation.

